# Collaborative study for the BINACLE assay for *in vitro* determination of botulinum neurotoxin activity

**DOI:** 10.64898/2026.09.01.748470

**Authors:** Birgit Kegel, Kay-Martin Hanschmann, Ursula Bonifas, Jolanta Klimek, Heike A. Behrensdorf-Nicol

**Affiliations:** Divisions of Veterinary Medicine, Paul-Ehrlich-Institut, Paul-Ehrlich-Straße 51-59, 63225 Langen, Germany; Divisions of Safety of Biomedicines and Diagnostics, Paul-Ehrlich-Institut, Paul-Ehrlich-Straße 51-59, 63225 Langen, Germany

**Keywords:** Botulinum neurotoxin, *in vitro*, potency testing, BINACLE (binding and cleavage) assay, collaborative study

## Abstract

The muscle-relaxing effects of botulinum neurotoxin (BoNT) serotypes A and B are widely utilized in clinical and aesthetic medicine. While the mouse bioassay remains the gold standard for evaluating the potency of pharmaceutical BoNT preparations, a non-animal alternative would be preferable to comply with European Directive 2010/63/EU. The Binding and Cleavage (BINACLE) assay, which measures BoNT activity based on the receptor-binding and proteolytic properties of the toxin, represents a promising alternative. Previous studies showed that this method allows reliable and sensitive activity determination.

Here, we describe a collaborative study involving ten international laboratories, which aimed to characterize the applicability of the BINACLE assay for the potency testing of pharmaceutical BoNT products. Our results demonstrate that the BINACLE assay enables reproducible relative potency determinations for BoNT/A, the most pharmaceutically relevant serotype, with variability within acceptable limits. Reproducible relative potency calculations were also demonstrated for BoNT/B, while the data highlighted opportunities for refinement of the BoNT/B BINACLE protocol to further enhance the robustness of the method.

Overall, this work confirms the utility of the BINACLE assay as a reliable and sensitive method for determining BoNT product potency, thereby substantiating its applicability as an alternative to the mouse bioassay.

## 1 Introduction

The botulinum neurotoxin (BoNT) serotypes A and B are used to treat a variety of disorders in clinical and aesthetic medicine. Due to their high toxicity, a precise potency determination of each pharmaceutical BoNT batch is important in compliance with the European Pharmacopoeia monographs 2113 and 2581 [1;2]. To date, the mouse bioassay, which is based on the determination of the median lethal dose (LD_50_), is considered as “gold standard” method for measuring BoNT activity. Accordingly, the activity units used to quantify the potency of the toxin products are based on the lethal doses determined in mice. However, this test has been reported to be associated with pronounced variability [3]. Besides, the mouse bioassay raises animal welfare concerns, as it requires high numbers of mice and causes severe distress for the test animals. For these reasons, the European Pharmacopoeia strongly encourages the development and validation of alternative methods for the potency determination of pharmaceutical BoNT products [1;2]. Therefore, our objective is to provide a freely available *in vitro* assay for the activity determination of BoNT/A and BoNT/B that can contribute to a replacement of the mouse bioassay.

BoNT molecules consist of two subunits connected by a disulfide bond: The heavy chain mediates the binding to receptors on target neurons and the uptake of the toxin into the cell. The light chain contains a proteolytic domain which cleaves SNARE (soluble N-ethylmaleimide-sensitive factor attachment protein receptor) proteins within the neurons, thereby inhibiting neurotransmitter release. The BoNT serotypes differ in their receptor and substrate preferences: BoNT/A uses ganglioside GT1b and the synaptic vesicle glycoprotein 2C (SV2C) as receptors for target cell binding [4;5]; thereafter, it specifically cleaves synaptosomal-associated protein-25 (SNAP-25) as substrate protein [6;7]. BoNT/B, in contrast, recognizes ganglioside GT1b and the protein synaptotagmin as receptors [8], while synaptobrevin, which is also called VAMP (vesicle-associated membrane protein), serves as substrate for its proteolytic activity [9].

The BINACLE (<u>bin</u>ding <u>a</u>nd <u>cle</u>avage) assay developed at the Paul-Ehrlich-Institut (PEI) measures BoNT activity *in vitro* based on the receptor binding function as well as the proteolytic activity of the toxin molecules. In the first step of this assay, the receptor molecules specific for the respective BoNT serotype are immobilized on a microplate and incubated with the test samples. Active toxin molecules bind to the receptors via their heavy chains and are then treated with a reducing agent in order to release and activate the light toxin chains. The supernatant containing the light chains is subsequently transferred to a second microplate containing an immobilized substrate protein, which is specifically cleaved by the active BoNT light chains. The resulting cleavage fragment is detected using a cleavage site-specific antibody. Depending on the receptor molecules and substrate proteins used, this test system can specifically measure the activity of either BoNT/A- or BoNT/B-containing samples [10;11]. In addition, BINACLE assays can also detect active tetanus neurotoxin when reagents specific for this toxin type are used [12]. In-house studies have shown that the BINACLE assays allow reliable and sensitive measurements of BoNT activity with a detection limit that is comparable to or even lower than the detection limit of the mouse bioassay [10;11]. In addition, a transferability study with four participants demonstrated that the BoNT BINACLE method can also be successfully performed in other laboratories [13]. To further examine the applicability of the BINACLE assay as an *in vitro* alternative to the LD_50_-based mouse bioassay for the activity and potency determination of BoNT products, we then initiated a collaborative study. In this study, identical toxin-containing sample solutions were measured in multiple BINACLE assays by ten international laboratories to provide an in-depth evaluation of the assay characteristics. The aim of this study was to characterize the applicability of the BINACLE assay under real-world conditions, thereby paving the way for the widespread use of the method.

## 2 Material and Methods

### 2.1 Material

The following materials, reagents and stock solutions for the BINACLE assays were provided by the Paul-Ehrlich-Institut (PEI) to all participants after prequalification at the PEI: Transparent MaxiSorp^®^ F96 microplates (Nunc A/S, Roskilde, Denmark, order no. 439454); a 1 mg/mL solution of ganglioside GT1b (Sigma-Aldrich, Taufkirchen, Germany, order no. G3767) in methanol; a 1 mg/mL solution of a recombinant protein representing amino acids 454-579 of human SV2C (toxologics GmbH, Hannover, Germany) in phosphate buffered saline (PBS); a 4 mg/mL solution of the peptide acetyl-GESQEDMFAKLKEKFFNEINKC (representing amino acids 40-60 of mouse synaptotagmin-2, synthesized by GeneCust, Dudelange, Luxembourg) in 8 mM ammonium hydrogen carbonate; a 15.6 µM solution of recombinant SNAP-25 (List Biological Laboratories, Campbell, CA, USA) in 7 mM HEPES, pH 7.4 with 0.5% lactose; a 175 µM solution of recombinant synaptobrevin (amino acids 1-97 of rat synaptobrevin-2 with an N-terminal histidine tag, manufactured by toxologics GmbH, Hannover, Germany, following a published protocol [14]) in 20 mM sodium acetate, pH 4.5, with 0.05% Triton^®^ X-100; affinity-purified polyclonal rabbit antibodies specifically recognizing either cleaved SNAP-25 (concentration: 1.1 µg/mL) or cleaved synaptobrevin (concentration: 5.0 µg/mL) in PBS with 1% BSA (contract-manufactured by Biotrend GmbH, Cologne, Germany, following published protocols [10;15]); biotin-conjugated goat-anti-rabbit IgG (Dianova GmbH, Hamburg, Germany, order no. 111-065-144, rehydrated in 50% glycerol according to the manufacturer’s instructions); peroxidase-conjugated streptavidin (Dianova GmbH, order no. 016-030-084, rehydrated in 50% glycerol according to the manufacturer’s instructions); a 0.5 M solution of Tris(2-carboxyethyl) phosphine hydrochloride (TCEP, from Sigma-Aldrich, order no. C4706) in water, pH adjusted to 6.8; 3,3′,5,5′-tetramethyl benzidine (TMB) solution (Life Technologies, Darmstadt, Germany, order no. 7588925); a 40 mg/ml solution of asolectin (Sigma-Aldrich, order no. 11145) in PBS; trimethylamine N-oxide dihydrate (TMAO, from Sigma-Aldrich, order no. 92277); protease-free bovine serum albumin (BSA, from Serva, Heidelberg, Germany, order no. 11926); Tween^®^ 20 (Sigma-Aldrich, order no. P7949); PBS solution (10x) without Ca^2+^, Mg^2+^ (Biochrom, Berlin, Germany, order no. L 1835); phosphate buffer without NaCl and KCl, pH 7.1 (prepared at the PEI); 100 mM 1,4-piperazinediethanesulfonic acid (PIPES), pH 6.4 (prepared at the PEI); 450 mM PIPES, pH 6.4 (prepared at the PEI). Chemicals that were considered non-critical, such as distilled water, ethanol, hydrochloric acid, sodium hydroxide and sulphuric acid, were purchased by each participant individually.

### 2.2 Sample and control solutions

Research-grade preparations of BoNT/A (specific toxicity: 2.6 x 10^8^ U/mg) and BoNT/B (specific toxicity: 1 x 10^8^ U/mg) without neurotoxin-associated proteins were purchased from Metabiologics Inc. (Madison, WI, USA). The pharmaceutical products IncobotulinumtoxinA (Xeomin^®^, Merz Pharma GmbH & Co. KGaA, Frankfurt, Germany) containing free BoNT/A, OnabotulinumtoxinA (Botox^®^, Allergan Pharmaceuticals, Westport, Ireland) containing complex-associated BoNT/A, and RimabotulinumtoxinB (Neurobloc^®^, Eisai Manufacturing Ltd, Hatfield, UK) containing complex-associated BoNT/B were purchased from a pharmacy. All products were used within their shelf-life period. Unless otherwise specified, all activity units (U) indicated in this article refer to the median lethal dose (LD_50_) for mice, as specified by the respective toxin manufacturer.

Lyophilized toxin products were reconstituted according to the manufacturer’s recommendations. Samples as well as positive and negative control solutions for the study were prepared at the PEI by diluting the respective toxin stock solutions in 9.6 mM salt-free phosphate buffer, pH 7.1 with 1% w/v BSA (for BoNT/A) or in PBS, pH 7.1 with 1% w/v BSA (for BoNT/B) to the concentrations indicated in Table 1. The samples were blinded and sent to the study participants in frozen aliquots together with the non-blinded positive and negative control solutions.

**Table 1:** Samples and control solutions prepared for the collaborative study.

|  | **Solutions for BoNT/A assays** | **Solutions for BoNT/B assays** |
| --- | --- | --- |
| **Positive Control** | Research-grade BoNT/A, 20 U/mL (corresponds to 76 pg/mL) | Research-grade BoNT/B, 0.67 U/mL (corresponds to 6.6 pg/mL) |
| **Negative Control** | No toxin (buffer only) | No toxin (buffer only) |
| **Sample 1** | No toxin (buffer only) | No toxin (buffer only) |
| **Sample 2** | Research-grade BoNT/A, 4.0 U/mL | Research-grade BoNT/B, 0.22 U/mL |
| **Sample 3** | Research-grade BoNT/A, 0.8 U/mL | Research-grade BoNT/B, 0.074 U/mL |
| **Sample 4** | Research-grade BoNT/A, 0.16 U/mL | Research-grade BoNT/B, 0.025 U/mL |
| **Sample 5** | OnabotulinumtoxinA, 4.0 U/mL | Research-grade BoNT/B, 0.008 U/mL |
| **Sample 6** | OnabotulinumtoxinA, 0.8 U/mL | RimabotulinumtoxinB, 2.0 U/mL |
| **Sample 7** | OnabotulinumtoxinA, 0.16 U/mL | RimabotulinumtoxinB, 0.67 U/mL |
| **Sample 8** | IncobotulinumtoxinA, 4.0 U/mL | RimabotulinumtoxinB, 0.22 U/mL |
| **Sample 9** | IncobotulinumtoxinA, 0.8 U/mL | RimabotulinumtoxinB, 0.074 U/mL |
| **Sample 10** | IncobotulinumtoxinA, 0.16 U/mL | RimabotulinumtoxinB, 0.025 U/mL |

### 2.3 BINACLE assays

General notes: The compositions of the buffers and working solutions used in the BINACLE assays for the activity determination of BoNT/A or BoNT/B are shown in Table 2. During incubation steps performed at 37°C or at room temperature, the microplates were gently agitated. Participants were free to wash the microplates either manually or by using an automated washer. Unless otherwise specified, each washing step consisted of four cycles, with 300 µL/well of Washing Buffer (Table 2) used in each cycle. When washing the Binding Plate before the addition of Reduction Buffer or the Cleavage Plate before proteolytic cleavage and before incubation with TMB, five wash cycles were applied for a particularly thorough removal of unbound material.

**Table 2:** Composition of buffers used in the BINACLE assays.

|  | **BINACLE assay for BoNT/A** | **BINACLE assay for BoNT/B** |
| --- | --- | --- |
| **Coating Solution for Binding Plate** | 5 µg/mL ganglioside GT1b and 10 µg/mL recombinant SV2C protein in PBS, pH 7.1 | 5 µg/mL ganglioside GT1b and 2.5 µg/mL synaptotagmin peptide in PBS, pH 7.1 |
| **Coating Solution for Cleavage Plate** | 100 nM recombinant SNAP-25 in PBS, pH 7.1 | 0.75 µM recombinant synaptobrevin in PBS, pH 7.1 |
| **Blocking Buffer for Binding Plate** | 1% w/v BSA in PBS, pH 7.1 |  |
| **Blocking Buffer for Cleavage Plate** | 1% w/v BSA and 100 µg/mL asolectin in PBS, pH 7.1 |  |
| **Washing Buffer** | 0.05% v/v Tween® 20 in PBS, pH 7.1 |  |
| **Binding Buffer** | 1% w/v BSA in phosphate buffer without NaCl and KCl, pH 7.1 | 1% w/v BSA in PBS, pH 7.1 |
| **Reduction Buffer** | 2.5 mM TCEP in phosphate buffer without NaCl and KCl, pH 7.1 | 2.5 mM TCEP in 100 mM PIPES, pH 6.4 |
| **Cleavage Buffer** | Prepared by dissolving 3 g TMAO in 6 mL 450 mM PIPES, pH 6.4 | Prepared by dissolving 4 g TMAO in 4 mL 100 mM PIPES, pH 6.4 |
| **Antibody Buffer** | 0.5% w/v BSA in PBS, pH 7.1 |  |

Each BINACLE assay was performed over a period of three consecutive days (as summarized in Figure 1):

**Figure 1:**
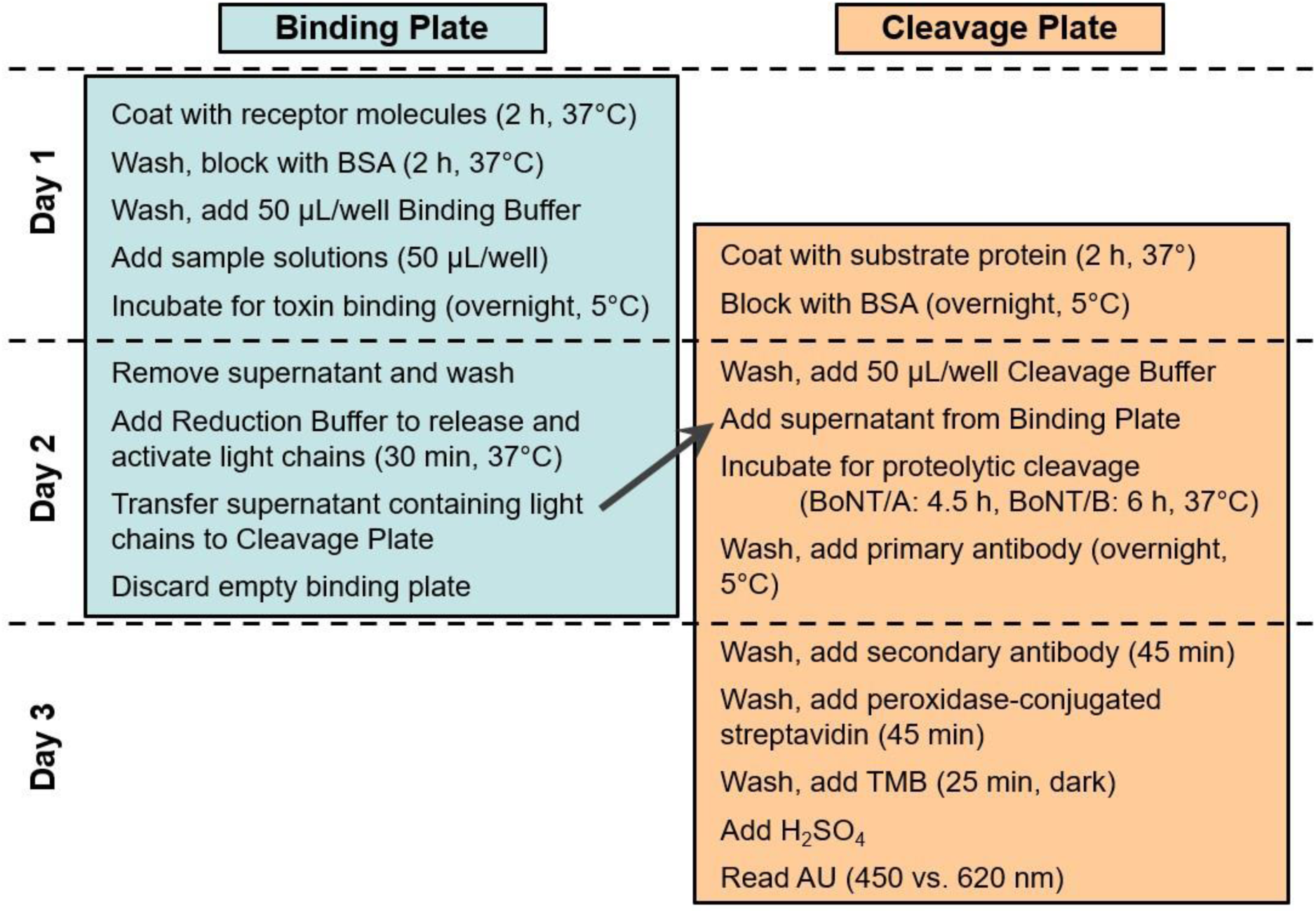
Key working steps of the BINACLE assays for determining the activity of BoNT. Unless otherwise indicated, incubations were performed at room temperature.

On day 1, the Binding Plate was prepared by coating a MaxiSorp^®^ F96 microplate with the specific receptor molecules for the corresponding BoNT serotype. For this, 100 µL/well Coating Solution for Binding Plate were added, and the plate was incubated for 2 h at 37°C. Control wells coated with PBS without receptor molecules were included in each assay to assess non-specific binding. After coating, the plate was washed, and unspecific protein binding sites were blocked by incubating the plate with 250 µL/well Blocking Buffer for Binding Plate for 2 h at 37°C. After this incubation, the Binding Plate was washed, and 50 µL of Binding Buffer were added to each well. Then 50 µL/well of the respective BoNT-containing samples as well as the positive and negative control solutions (Table 1) were added to the Binding Plate in multiple replicates each, following a predefined pipetting scheme (Figure 2). The plate was then incubated overnight at 5°C to allow toxin binding to the immobilized receptors. In parallel, the Cleavage Plate was prepared by coating a fresh MaxiSorp microplate with substrate proteins. For this, 100 µL Coating Solution for Cleavage Plate were added to each well, and the plate was incubated for 2 h at 37°C. Subsequently, unspecific protein binding sites were blocked by incubating the plate overnight at 5°C with 250 µL/well Blocking Buffer for Cleavage Plate.

**Figure 2:**
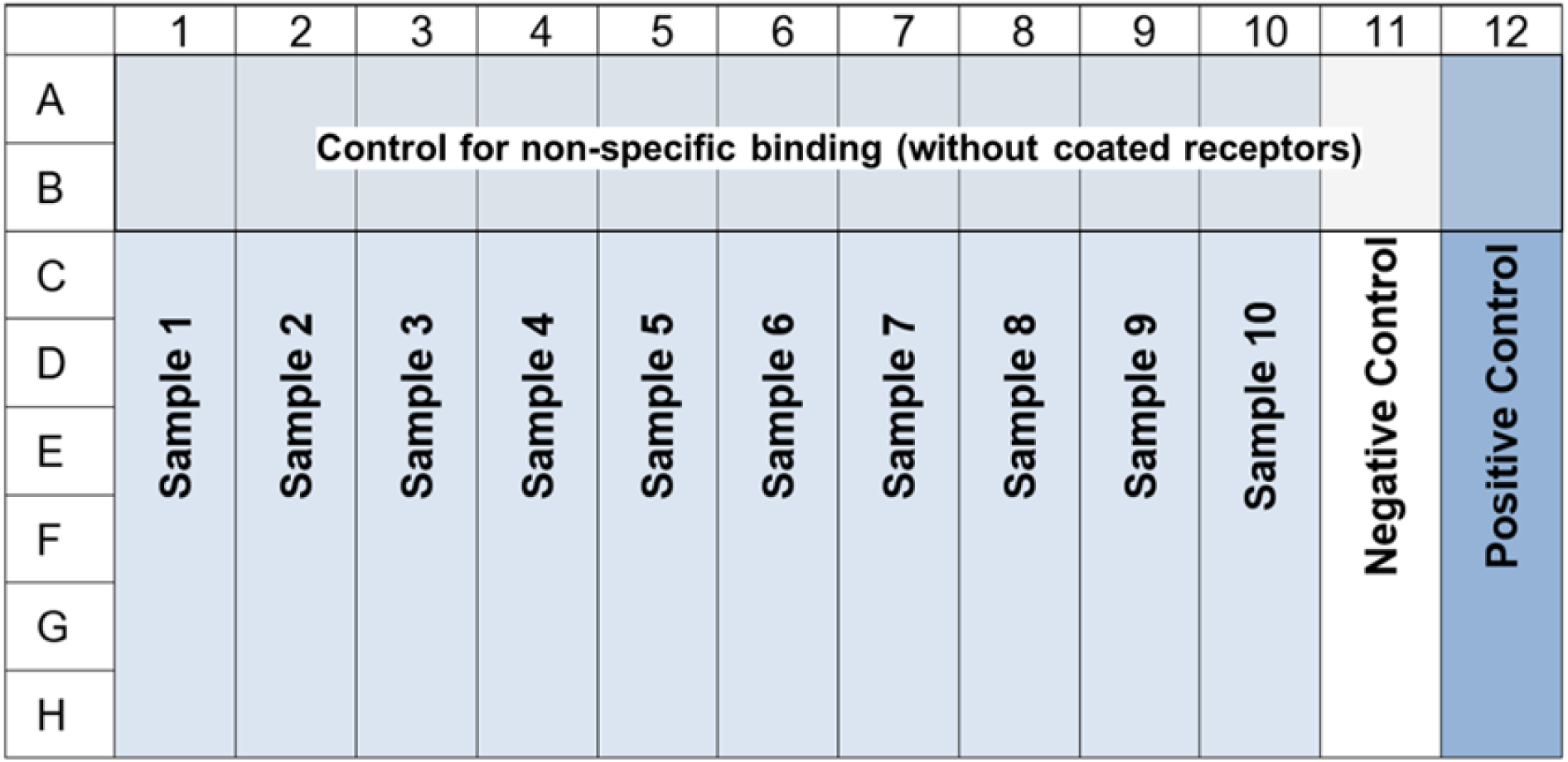
Plate layout for the BINACLE assays. performed in the collaborative study. The composition of the samples and the negative and positive control solutions used in the assays for activity determination of BoNT/A and BoNT/B is shown in Table 1. Each sample or control solution was added to six receptor-coated microplate wells (in rows C to H of the plate) and two wells without receptor molecules (in rows A and B), which served as control to determine the background signals induced by non-specific binding.

On day 2, the Binding Plate containing the bound toxin molecules was washed. In BoNT/B assays, the Washing Buffer described in Table 2 was used for this step, whereas in BoNT/A assays, phosphate buffer without NaCl and KCl, pH 7.1 with 0.05% Tween^®^ 20 was used instead. Then 100 µL/well of Reduction Buffer were added, and the Binding Plate was incubated for 30 min at 37°C to induce the release and activation of the light toxin chains. In parallel to this incubation, the Cleavage Plate containing the immobilized substrate proteins was washed, and 50 µL Cleavage Buffer were then added to each well of the Cleavage Plate. Afterwards, the supernatants of the reduction step were transferred from the wells of the Binding Plate to the wells of the Cleavage Plate. The empty Binding Plate was discarded, and the Cleavage Plate was incubated at 37°C for 4.5 h (in BoNT/A assays) or 6 h (in BoNT/B assays) to allow the light chain-induced cleavage of the substrate proteins. Following this cleavage incubation, the Cleavage Plate was washed. Then 100 µL/well of the antibody specifically recognizing cleaved SNAP-25, diluted 1:1,000 in Antibody Buffer (in BoNT/A assays), or the antibody specifically recognizing cleaved synaptobrevin, diluted 1:4,000 in Antibody Buffer (in BoNT/B assays), were added to the plate for an overnight incubation at 5°C. On day 3, the Cleavage Plate was washed, and 100 µL of biotin-conjugated goat-anti-rabbit IgG diluted 1:2,500 in Antibody Buffer were added to each well and incubated at room temperature for 45 min. Then the plate was washed again and incubated with 100 µL/well peroxidase-conjugated streptavidin diluted 1:8,000 in Antibody Buffer for 45 min at room temperature. Afterwards, the plate was washed and incubated with 100 µL/well TMB solution for 25 min at room temperature in the dark without agitation. Finally, the color reaction was stopped by adding 50 µL/well of 1 M H_2_SO_4_, and the absorbance units (AU) for each well were measured in a microplate reader at 450 nm against 620 nm as reference wavelength.

### 2.4. Study design

Before the start of the experimental phase, the participants received detailed instructions for the performance of the BINACLE assays, and they were offered the opportunity to attend an optional BINACLE training course at the PEI. During the experimental phase, each participant was requested to perform three independent BINACLE assays for BoNT/A and three independent BINACLE assays for BoNT/B using the reagents and the BoNT-containing sample and control solutions provided by the PEI (see chapters 2.1 and 2.2). As some of the provided reagents had a limited shelf life, the participants were asked to carry out all tests within six months of receiving them.

Each assay was to be performed in accordance with the instructions given in the study protocol. After each test, participants were asked to check whether the following indicators of successful performance had been met:

- The signals obtained for the positive control should clearly exceed the signals obtained for the negative control.
- Background signals measured for the negative control solution and in control wells that were not coated with receptor molecules should be well below 1.0 AU.

After completion of the experiments, the participants sent the results of their photometric measurements to the PEI for statistical analysis. Participants were also asked to report any unexpected observations or deviations from the test protocol that occurred while performing the assays.

### 2.5 Follow-up tests for optimizing the washing conditions for BoNT/B

To further investigate unforeseen effects observed during the analysis of the BoNT/B data obtained during the study, follow-up experiments were conducted to assess the impact of the washing conditions on the synaptobrevin cleavage in the BoNT/B BINACLE assay. Three distinct washing procedures were compared:

(1.) Automated washing: All steps were performed as outlined in chapter 2.3, utilizing an automated washer for microplate washing.

(2.) Manual washing: All steps adhered to the description in chapter 2.3, with manual washing of the plates.

(3.) Modified manual washing: Washing was performed manually, and the following deviation from the protocol described in chapter 2.3 was applied: When washing the Binding Plate before the addition of Reduction Buffer and when washing the Cleavage Plate before the addition of Cleavage Buffer, four cycles of Washing Buffer (Table 2) were applied, followed by one final cycle using 100 mM PIPES (pH 6.4) without detergent. This detergent-free buffer was chosen because its composition closely matched that of the PIPES-based buffers used in the subsequent incubation steps (Reduction Buffer and Cleavage Buffer).

### 2.6 Statistical methods

During assay performance, all toxin-containing sample and control solutions were mixed with an equal volume of Binding Buffer (see chapter 2.3). Consequently, the final toxin concentrations in the microplate wells were half those of the initial solutions described in Table 1. As the data analysis was performed based on these final toxin concentrations, all concentrations indicated in the “Results” section of this article refer to the solutions in the microplate.

For the statistical data analysis, the participants were randomly assigned the laboratory codes “A” to “J”. Descriptive statistics (e.g. mean, standard deviation) were calculated for each sample and positive control solution from the AU measured in the respective wells. The calculation of relative potency (RP) values was performed with CombiStats software, version 6.1 [16] using a four-parameter sigmoid dose-response-curve model as described in the European Pharmacopoeia [17]. In one case, an additional asymmetry factor was included in the curve fitting (five-parameter logistic dose-response curve).

The sigmoid curves were checked for parallelism and linearity (linearity for the logit-transformed curve data) by means of an F-test testing for significant deviations from parallelism and/or linearity. For parallelism, acceptance criterion was a P value for non-parallelism above 0.05 or, in case the P value was at least above 0.01, a 90% confidence interval for the ratio of slopes in between acceptance limits of 0.63 – 1.60. Linearity was considered acceptable, if the P value for non-linearity was above 0.05. P values for non-linearity above 0.01 were exceptionally accepted in cases where at least half of the sigmoid curve shape was supported by the data and parallelism of the curves was evident.

As no certified BoNT standard preparations were available, the RP values for the pharmaceutical BoNT products were calculated relative to the research-grade BoNT preparations, which served as internal reference samples in the study. Based on the manufacturer’s activity specifications, the following potencies were assigned to the respective 1 mg/mL stock solutions: 2.6 × 10^8^ U/mL for research-grade BoNT/A, 1.0 × 10^8^ U/mL for research-grade BoNT/B.

Combined RP results for each laboratory and BoNT product were obtained by combining the valid individual assays with CombiStats. In case of sufficient homogeneity (defined as a p value for the homogeneity test of ≥0.100), the weighted combination was used; otherwise, the semi-weighted combination was used.

All other statistical analyses were performed with SAS^®^/STAT software, version 9.4, SAS System for Windows [18]. The coefficient of variation was used for description of relative variability of the measurements. The influence of relevant factors, such as the laboratory, on the inter-assay precision (intermediate precision) and the intra-assay precision (repeatability) was evaluated by means of a mixed linear model (an analysis of variance, ANOVA, using random factors). This method uses (restricted) maximum likelihood estimates, which may lead to a small difference between the estimated variance and the usual variance estimator.

The inter-assay precision was described by the standard deviation and the coefficient of variation, derived from the total variation (using the overall mean estimator). For the intra-assay precision, the residual variance was used. The measurement uncertainty was then described as the estimated total variance from the ANOVA, also denoted as coefficient of variation.

## 3 Results

### 3.1 Test performance and data included in statistical analysis

Ten international participants, including medicines control laboratories, national health authorities, pharmaceutical companies, and research laboratories, took part in the collaborative study. Each participant performed three consecutive BINACLE assays for BoNT/A and three consecutive assays for BoNT/B.

In the BINACLE protocol provided to the participants, a clearly visible difference between the positive and negative control signals had been defined as a key indicator for successful test performance. This criterion was not met in one BoNT/A assay conducted by laboratory D and in one BoNT/B assay conducted by laboratory G. In these assays, the positive control solutions yielded only low signals which did not clearly exceed the negative control signals, suggesting that experimental errors may have occurred. Therefore, each of these two laboratories performed an additional repeat test, the results of which were included in the data analysis instead of those of the original non-compliant test. Consequently, data from 30 tests (ten laboratories, three tests each) for BoNT/A and 30 tests for BoNT/B (ten laboratories, three tests each) were subjected to statistical evaluation.

All tests were performed in close compliance with the protocol, only minor deviations from the protocol instructions were reported by the participants. For example, in some cases, the shaking conditions during microplate incubations or the centrifugation conditions for the sample vials had to be adapted to the specifications of the device types that were available in the respective laboratories. These deviations were not expected to exert any influence on the resulting assay signals.

Six single microplate wells (four wells from BoNT/A tests and two wells from BoNT/B tests) showing unreasonably high or unreasonably low signals were removed from data analysis because the participants had reported pipetting errors or washing problems for these wells.

### 3.2 Signals measured in the BINACLE assays

In the blank wells containing buffer without toxin as well as in wells without coated receptor molecules that served as control for non-specific binding, mean signals below 0.3 AU were obtained by all laboratories in the BoNT/A and the BoNT/B tests (Table 3). Laboratory J reported the highest background signals (between 0.2 and 0.3 AU), whereas laboratory E measured the lowest background (between 0.04 and 0.06 AU). These background values are within the usual range for BINACLE measurements.

**Table 3:** Control signals measured in the BINACLE assays. For each laboratory, the mean AU measured in blank wells without toxin (N=36, resulting from 3 assays with 12 replicate wells each) and in control wells without receptors (N=72, resulting from 3 assays with 24 replicate wells each) are shown with the corresponding standard deviation (SD).

| Lab | BoNT/A-BINACLE assays |  |  |  | BoNT/B-BINACLE assays |  |  |  |
| --- | --- | --- | --- | --- | --- | --- | --- | --- |
|  | Blank |  | No receptor |  | Blank |  | No receptor |  |
|  | Mean | SD | Mean | SD | Mean | SD | Mean | SD |
| A | 0.077 | 0.010 | 0.079 | 0.008 | 0.125 | 0.014 | 0.118 | 0.009 |
| B | 0.128 | 0.037 | 0.152 | 0.046 | 0.170 | 0.053 | 0.170 | 0.067 |
| C | 0.111 | 0.042 | 0.110 | 0.043 | 0.082 | 0.026 | 0.074 | 0.009 |
| D | 0.092 | 0.016 | 0.095 | 0.017 | 0.077 | 0.017 | 0.074 | 0.014 |
| E | 0.052 | 0.004 | 0.055 | 0.005 | 0.042 | 0.003 | 0.043 | 0.005 |
| F | 0.163 | 0.046 | 0.168 | 0.074 | 0.125 | 0.051 | 0.131 | 0.040 |
| G | 0.159 | 0.027 | 0.182 | 0.062 | 0.110 | 0.011 | 0.110 | 0.010 |
| H | 0.155 | 0.039 | 0.174<sup>§</sup> | 0.054<sup>§</sup> | 0.164 | 0.031 | 0.169 | 0.050 |
| I | 0.120 | 0.045 | 0.125 | 0.044 | 0.103 | 0.017 | 0.103 | 0.020 |
| J | 0.226 | 0.023 | 0.252 | 0.016 | 0.223 | 0.037 | 0.217 | 0.031 |

<sup>§</sup> Calculated from N=70 values, as two corresponding wells had been removed from analysis.

When measuring toxin-containing solutions in the BoNT/A BINACLE tests, all participants observed a clear increase of the measured signals with increasing toxin concentrations. This dose-dependent increase was seen for all tested BoNT/A preparations, i.e. for research-grade BoNT/A, OnabotulinumtoxinA and IncobotulinumtoxinA (Figure 3A). Already for the wells with the lowest BoNT/A concentrations (0.08 U/mL, which corresponds to 0.3 pg/mL for the research-grade toxin preparation), the measured signals clearly exceeded the blank signal in most laboratories.

**Figure 3.**
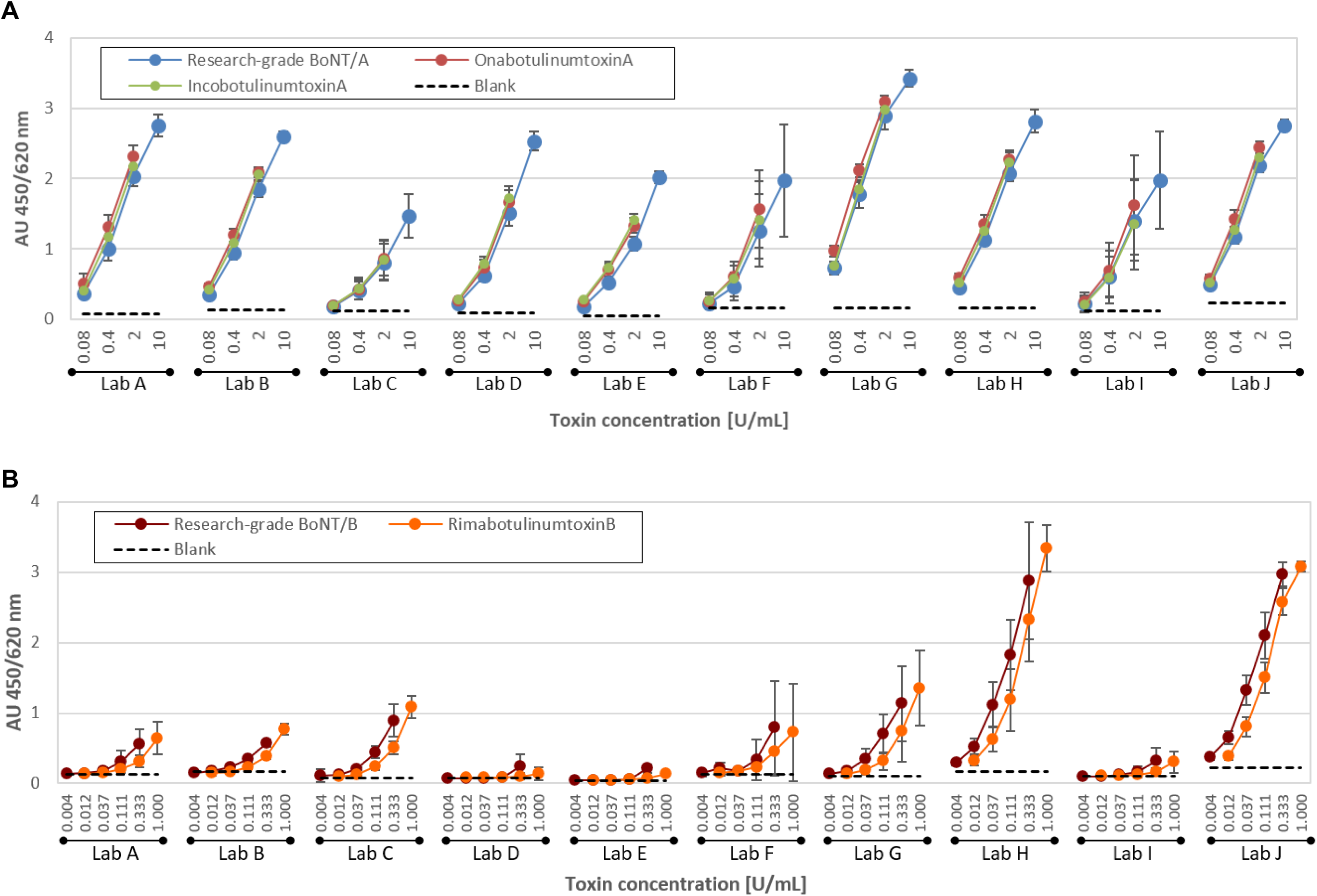
Mean signals obtained by the individual laboratories in the BoNT/A-BINACLE assays (A) for research-grade BoNT/A (blue symbols), OnabotulinumtoxinA (red symbols) and IncobotulinumtoxinA (green symbols), or in the BoNT/B BINACLE assays (B) for research-grade BoNT/B (brown symbols) and RimabotulinumtoxinB (orange symbols). The x-axes represent the final toxin concentrations on the microplate in U/mL (based on the unit specifications of the respective manufacturers). The y-axes represent the signals (AU) measured at 450 vs. 620 nm. Unless stated otherwise, each symbol represents the mean signal from N=18 measurements (three tests, six replicate wells each). Calculations for the following data points were based on N=17 wells, as one of the corresponding replicate wells was excluded from data analysis (see chapter 3.1): laboratory C, 2 U/mL IncobotulinumtoxinA; laboratory H, 0.4 U/mL IncobotulinumtoxinA; laboratory E, 0.33 U/mL research-grade BoNT/B; laboratory H, 1 U/mL RimabotulinumtoxinB. Standard deviations are indicated as error bars. The dotted lines represent the corresponding mean blank values.

For solutions containing 10 U/mL of research-grade BoNT/A (corresponding to a concentration of 38 pg/mL), the mean net signals measured by 8 out of 10 participating laboratories fell within the range of 1.8 to 2.7 AU (Figure 4A). Only laboratory C reported lower net signals with a mean value of 1.36 AU, whereas laboratory G measured the highest BoNT/A-induced signals among all participants with a mean value of 3.26 AU.

**Figure 4:**
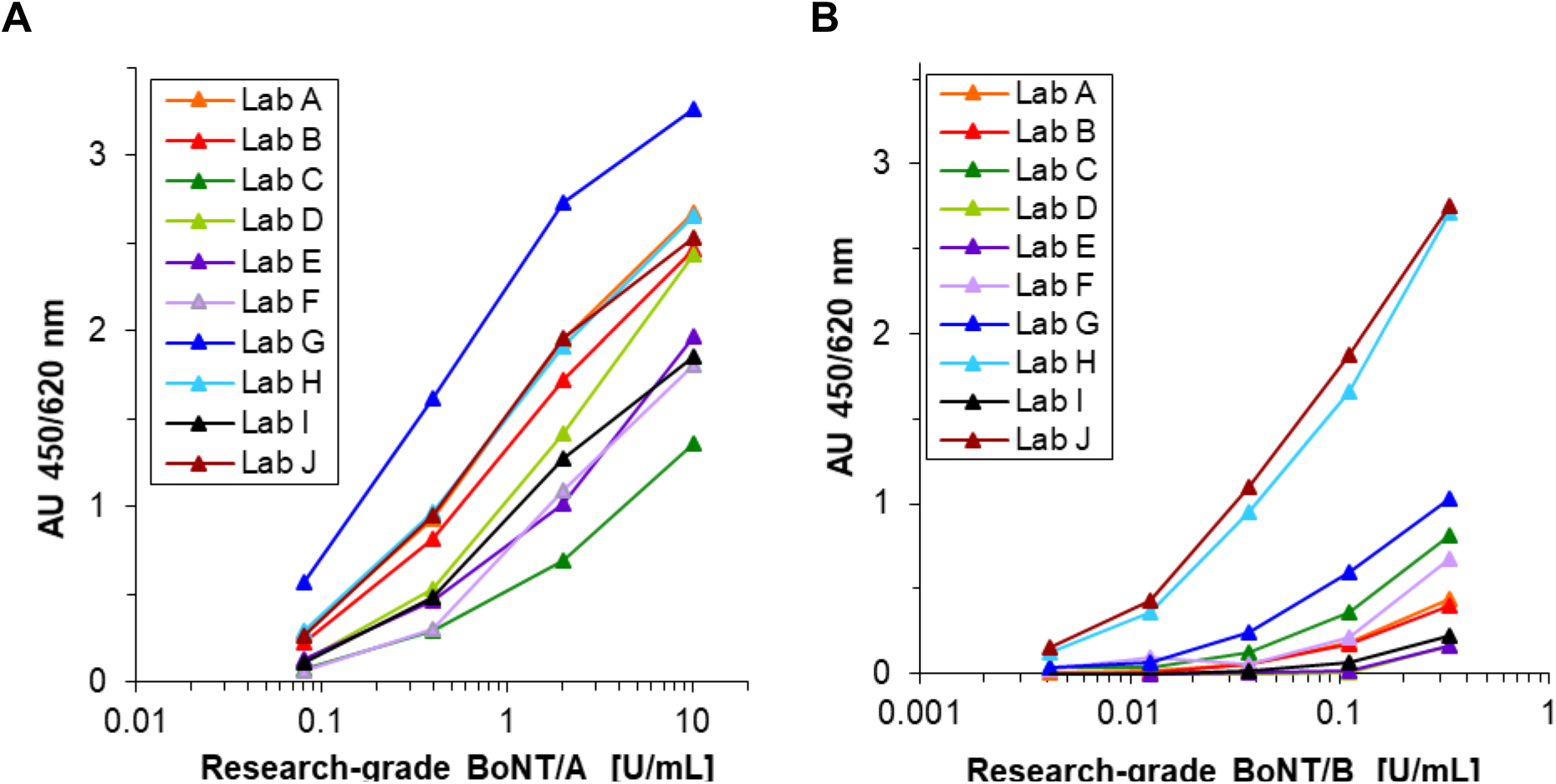
Comparison of mean net signals obtained by the individual participants for research-grade BoNT/A (A) and research-grade BoNT/B (B). The x-axes represent the final toxin concentrations on the microplate in U/mL (based on the unit specifications of the respective manufacturers). The y-axes represent the mean of the measured signals from N=18 microplate wells (3 tests, 6 replicates each) after subtraction of the corresponding blank signal without toxin. For the 0.33 /mL BoNT/B concentration in laboratory E, the calculation was based on N=17 wells, as one well had been excluded from data analysis (see chapter 3.1). Each laboratory is represented by a different color (see color legend in the graphs). In diagram (B), the curves for laboratories A and D are not clearly visible, as they strongly overlap with the curves for laboratories B and E, respectively. Similarly, in diagram (A), the curves for laboratories A, H and J partially mask each other.

Variability between the individual test runs was quite low in most laboratories, as illustrated by small error bars in Figure 3A. Only the signals measured by laboratories F and I show larger error bars, which indicates that these laboratories experienced a comparatively high inter-assay variability.

In the BoNT/B BINACLE assays performed during the study, dose-dependent toxin signals were also obtained in all laboratories (Figure 3B). Thus, measurements of BoNT/B activity were successfully achieved by all participants, in principle. However, large differences were found between the signal levels measured by the individual laboratories (Figure 3B; Figure 4B):

Participants H and J obtained comparatively high toxin signals. In these laboratories, even the lowest tested BoNT/B concentration of 0.004 U/mL induced signals that clearly exceeded the blank value, and the net signals obtained for 0.33 U/mL research-grade BoNT/B (which corresponds to a toxin concentration of 3.3 pg/mL) were close to 2.7 AU.

The other eight participants, in contrast, reported much lower toxin signals: In these laboratories, the AU values induced by low BoNT/B concentrations (i.e. 0.004 U/mL or 0.012 U/mL) barely exceeded the blank value, and the net signals measured for a 0.33 U/mL solution of research-grade BoNT/B were also much lower than those reported by laboratories H and J, with values ranging from 0.1 to 1.1 AU.

In addition to the high inter-laboratory variability observed for the BoNT/B-induced signals, some participants (e.g. laboratories F, G and H) also reported pronounced variability between the signal levels measured in their individual test runs, as illustrated by the large error bars in Figure 3B.

### 3.3 Relative potency values

From the data obtained in the BoNT/A tests, potency values were calculated for OnabotulinumtoxinA and IncobotulinumtoxinA relative to the research-grade BoNT/A, which served as internal reference toxin. All data sets generated during the study permitted the calculation of valid relative potencies (RP) (Table 4), with the following exceptions: Although the BoNT/A signals measured by laboratory E showed clear dose-dependent responses (Figure 3A), none of their three data sets allowed RP calculations for OnabotulinumtoxinA due to deviations from the predefined acceptance criteria for linearity or parallelism, and one data set also did not allow valid potency estimates for IncobotulinumtoxinA. Besides, test number 3 performed by laboratory I gave invalid results for both BoNT/A products.

**Table 4:**
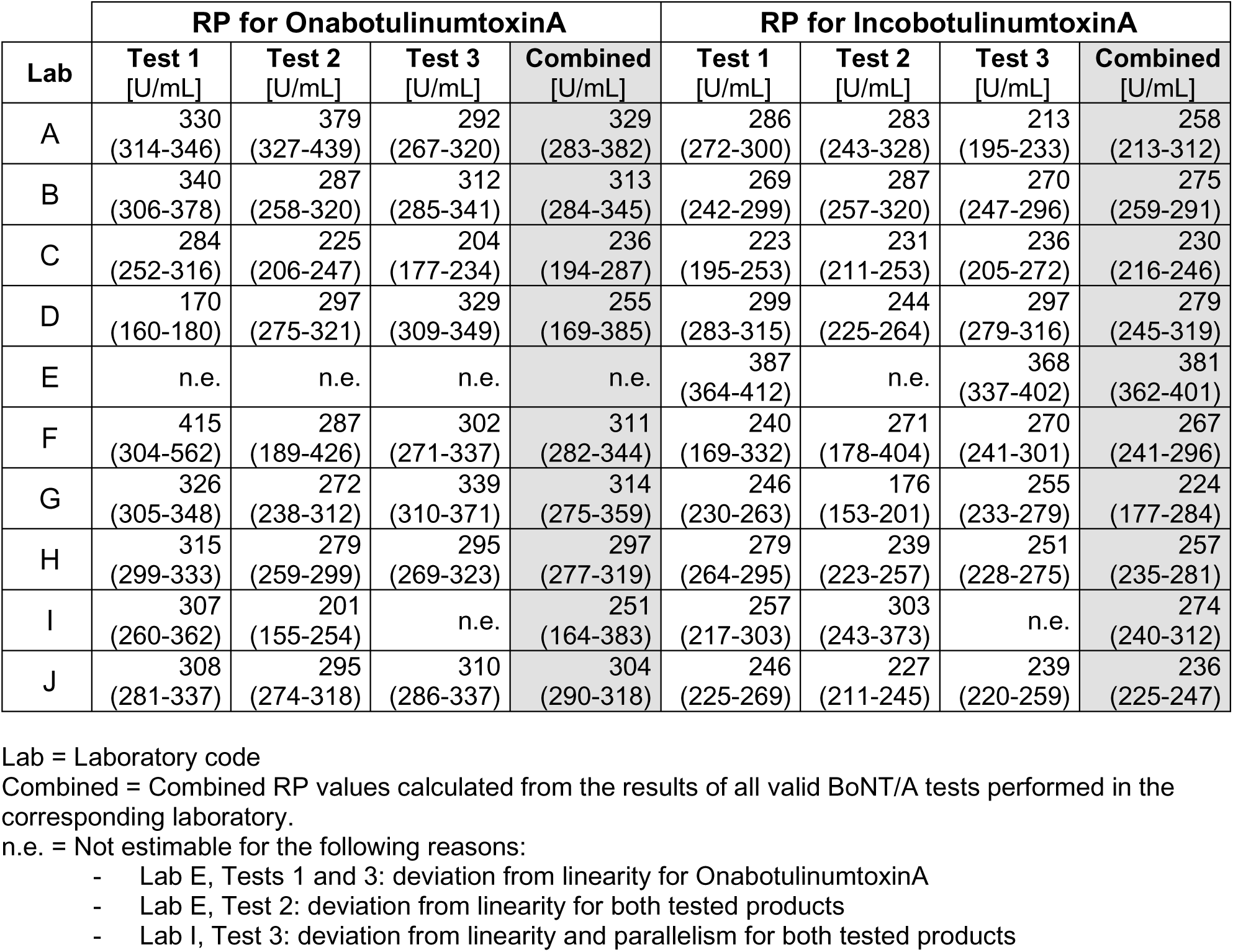
Relative potency estimates obtained for the BoNT/A products with 95% confidence intervals.

Combined RP values were calculated for each participant based on all valid data sets generated in the corresponding laboratory (Table 4). As illustrated by Figure 5, the combined RP estimates of the individual laboratories are quite close to each other: For OnabotulinumtoxinA, the combined RP estimates obtained in the study ranged from 236 U/mL (in laboratory C) to 329 U/mL (in laboratory A). For IncobotulinumtoxinA, the RP estimates were in the range between 220 and 280 U/mL for nine participants, whereas laboratory E obtained a higher RP estimate of 380 U/mL.

**Figure 5:**
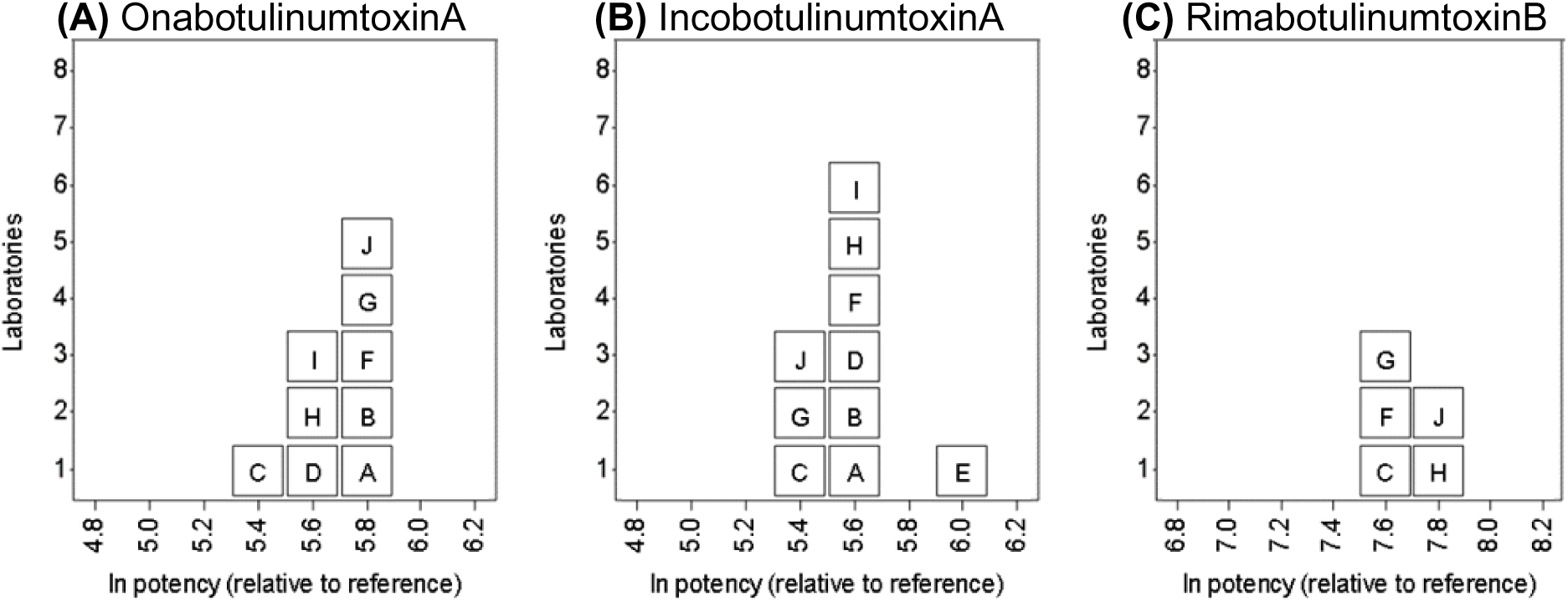
Graphical illustration of the RP values that were calculated for the tested pharmaceutical BoNT products based on the data of each participant. Each box represents the relative potency value that was calculated for OnabotulinumtoxinA (A), IncobotulinumtoxinA (B) and RimabotulinumtoxinB (C) from the combined data of all valid BINACLE tests performed in the respective laboratory. The letter in each box represents the laboratory code. The x-axes of the graphs indicate the natural logarithm (ln) of the respective potency values. The y-axis indicates the number of laboratories that obtained potency values in the corresponding range. Laboratory E obtained no valid potency estimate for OnabotulinumtoxinA; laboratories A, B, D, E, and I obtained no valid potency estimates for RimabotulinumtoxinB.

The measurement uncertainty of the RP estimates obtained for the BoNT/A products during the study did not exceed 20% (Table 5). Both the variability between the laboratories and the repeatability within one laboratory contributed to this overall measurement uncertainty to a roughly similar extent.

**Table 5:** Variability of RP estimates.

| Sample | Intercept | Covariance Parameter | Estimated Variance | Estimated Standard Deviation | Coefficient of Variation | Variability |
| --- | --- | --- | --- | --- | --- | --- |
| Onabotulinum-toxinA | 5.67 | Lab | 0.004 | 0.066 | 7% | Inter-assay |
|  |  | Residual | 0.033 | 0.183 | 18% | Intra-assay |
|  |  | Total variability | 0.038 | 0.195 | 20% | Overall |
| Incobotulinum-toxinA | 5.57 | Lab | 0.016 | 0.125 | 13% | Inter-assay |
|  |  | Residual | 0.012 | 0.109 | 11% | Intra-assay |
|  |  | Total variability | 0.028 | 0.166 | 17% | Overall |
| Rimabotulinum-toxinB | 7.71 | Lab | 0.011 | 0.106 | 11% | Inter-assay |
|  |  | Residual | 0.032 | 0.180 | 18% | Intra-assay |
|  |  | Total variability | 0.044 | 0.209 | 21% | Overall |

Intercept = mean RP estimate from least squares model, indicated as natural logarithm; inter-assay = variability between the laboratories; intra-assay = repeatability within one laboratory; overall = measurement uncertainty

In the BoNT/B tests, the research-grade BoNT/B preparation was used as internal reference toxin for calculating the RP of RimabotulinumtoxinB. Data analysis revealed that several participants encountered considerable difficulties in generating valid potency data for BoNT/B: We found that laboratories H and J, which had measured comparatively high signals for BoNT/B (see Figure 4B), obtained valid data sets from each of their tests (Table 6). In contrast, participants A, B, D, E and I, who reported much lower overall signal intensities for BoNT/B, did not obtain any valid data sets. Additionally, some data sets from laboratories C, F and G, which obtained signal intensities in the intermediate range, also failed to meet the predefined acceptance criteria. Overall, only 10 out of 30 BoNT/B BINACLE assays conducted during the study permitted valid RP calculations (Table 6).

**Table 6:** Relative potency estimates obtained for the BoNT/B product with 95% confidence intervals.

| Lab | RP for RimabotulinumtoxinB |  |  |  |
| --- | --- | --- | --- | --- |
|  | Test 1<br>[U/mL] | Test 2<br>[U/mL] | Test 3<br>[U/mL] | Combined<br>[U/mL] |
| C | n.e. | 1905<br>(1705-2129) | n.e. | 1905<br>(1705-2129) |
| F | 2207<br>(1778-2755) | n.e.<sup>§</sup> | n.e.<sup>§</sup> | 2207<br>(1778-2755) |
| G | 1537<br>(1274-1851) | n.e. | 2159<br>(1831-2553) | 1827<br>(1300-2567) |
| H | 3406<br>(3060-3808) | 2280<br>(1976-2634) | 2349<br>(2128-2593) | 2637<br>(2037-3414) |
| J | 2641<br>(2304-3030) | 2170<br>(1862-2529) | 2395<br>(2086-2753) | 2411<br>(2221-2618) |

Lab = Laboratory code. Laboratories A, B, D, E and I are not included in the table, as none of their BoNT/B tests allowed valid RP estimates.
Combined = Combined RP values calculated from the results of all valid BoNT/B tests performed in the corresponding laboratory.
n.e. = Not estimable due to deviations from linearity and/or parallelism.
§ = In these tests, no convergence was reached for the model fit, as the data points only covered the lower range of the sigmoid curve and showed high variability between replicates. The tests were rated as invalid.

Despite these challenges, when calculating the combined RP values for RimabotulinumtoxinB based on all valid test runs of each laboratory, we observed that the potency values obtained by the individual participants were within a fairly narrow range (Figure 5C), extending from 1827 U/mL (laboratory G) to 2637 U/mL (laboratory H) (Table 6). The measurement uncertainty associated with these RP estimates for RimabotulinumtoxinB was determined to be 21% (Table 5), indicating a moderate level of variability.

### 3.4 Follow-up experiments to optimize the BoNT/B-BINACLE assay

Figure 4B illustrates that the signal intensities obtained for the BoNT/B samples differed considerably between participants. An investigation into the potential causes of these differences revealed that the conditions employed for microplate washing were likely a contributing factor. The study protocol did not specify whether participants should wash the microplates by hand or use an automated plate washer. A review of the methods employed by the participants showed a notable correlation between washing methods and signal intensities: Among the four laboratories with the highest BoNT/B-signals, three laboratories (laboratories C, G and J) utilized an automated washer, and one (laboratory H) employed a hybrid approach, where washing buffer was added manually, but aspiration was performed using a washer. In contrast, the laboratories that reported the lowest BoNT/B signals (laboratories D, E, and I) performed washing by hand. This observation suggests that the use of automated washing methods may contribute to higher signal intensities in the BoNT/B BINACLE assay.

Follow-up experiments conducted at the PEI confirmed that the use of a microplate washer in the BoNT/B BINACLE assay yields substantially higher toxin signals compared to manual washing (Figure 6), thereby confirming the impact of washing conditions on the resulting data.

**Figure 6:**
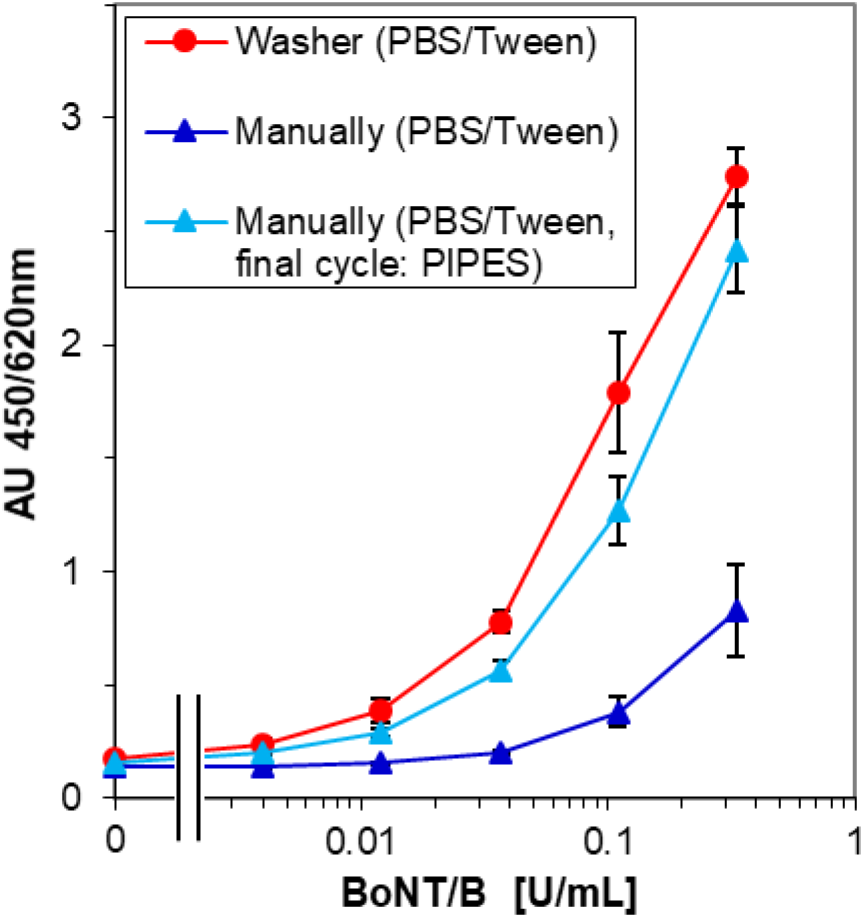
BINACLE signals obtained for research-grade BoNT/B using different washing conditions. The x-axis represents the final toxin concentration on the microplate in U/mL, the y-axis represents the signals (AU) measured at 450 vs. 620 nm. The washing steps were either performed with an automated microplate washer using PBS/0.05% Tween^®^ 20 as washing buffer (red circles), or manual washing was applied (light blue and dark blue triangles). For manual washing, two variants were compared: In one variant (dark blue triangles), PBS/0.05% Tween^®^ 20 was used for all washing steps. In the other variant (light blue triangles), PBS/0.05% Tween^®^ 20 was also used in most washing steps, but in the final washing cycles immediately before the toxin reduction and the synaptobrevin cleavage steps, it was replaced with a 100 mM PIPES buffer without detergent (see chapter 2.5). Each symbol represents the mean of n=9 measured values, derived from three independent tests with three replicate wells each. Standard deviations are indicated as error bars.

A relevant difference between manual and automated washing procedures may concern the aspiration step: While most microplate washers remove liquid from the wells stringently, manual washing procedures can result in a less thorough buffer removal, in our experience. Based on previous observations [19], we hypothesized that an elevated amount of washing buffer (PBS/0.05% Tween^®^ 20) remaining in the plate after manual washing could affect the synaptobrevin cleavage step in the BoNT/B BINACLE assay due to its detergent content. Consequently, we investigated whether removal of Tween^®^ 20 during the washing cycles preceding the proteolytic reaction would improve the resulting BoNT/B signals. Indeed, our experiments confirmed that, when applying manual microplate washing, much higher toxin signals were obtained when a detergent-free PIPES buffer was used instead of the regular washing buffer in the final washing cycles immediately prior to the toxin reduction step and the synaptobrevin cleavage step (Figure 6). This observation supports our hypothesis that elevated amounts of residual Tween^®^ 20 can compromise the proteolytic activity of BoNT/B.

In contrast, we did not find any evidence for a pronounced impact of the washing conditions on the toxin signals in the BoNT/A BINACLE assay (Figure 4A), as no systematic differences in BoNT/A signal intensities were found between study participants who had used a washer (e.g., laboratories C or J) and those who had washed the plates manually (e.g., laboratories D or E).

## 4 Discussion

To characterize the applicability of the BINACLE assay for activity determination of the botulinum neurotoxin serotypes A and B, identical toxin samples were tested in BINACLE assays by ten international laboratories within the framework of a collaborative study.

The BINACLE assays for the activity determination of serotype BoNT/A yielded the following overall results:

All ten laboratories successfully performed the BINACLE assays, and in each test run, a toxin concentration-dependent increase of the assay signals was observed, which illustrates the method’s robustness. The background signals in control wells without receptor or in wells without added toxin were in an acceptable range (i.e. below 0.3). In most tests performed during the study, already samples containing 0.08 U/mL BoNT/A induced signals that clearly exceeded the blank value. As each manufacturer-defined BoNT activity unit represents one mouse LD_50_, these results confirm that the BoNT/A BINACLE assay has a lower detection limit than the mouse bioassay. For samples containing 10 U/mL research-grade BoNT/A, high signals close to or above 2.0 AU were measured in almost all laboratories.

The overall signal intensities varied to some extent between the study participants. As the potency of the tested toxin products are usually determined relative to a reference toxin [1] rather than being calculated directly from the measured signals, such inter-laboratory variations in signal levels are not expected to impede reliable BoNT potency determinations. Our results emphasize the importance of including a well-characterized BoNT preparation as reference toxin in each test, against which the activity of the tested toxins can be calculated.

The RP values calculated for the pharmaceutical products IncobotulinumtoxinA and OnabotulinumtoxinA from the test results obtained in the individual laboratories were all within a fairly narrow range, whereby the total variability of the RP estimates did not exceed 20%. These findings underline the reliability of the BINACLE assay, as this level of variability falls within the range that is typically observed for multi-step immunochemical methods.

Only one of the ten participants had severe difficulties generating valid RP estimates for BoNT/A due to deviations from the acceptance criteria (parallelism or linearity). This failure to meet the criteria was somewhat unexpected, because at first glance, the dose-response curves generated in this laboratory were not noticeably different from those of the other laboratories. This difficulty in generating valid data may be partly due to the study design, which only comprised a small number of BoNT/A dilutions. Including more toxin dilutions in future applications of the BINACLE method to provide more comprehensive coverage of the dose-response curve would facilitate the reliable evaluation of assay signals and compliance with the acceptance criteria.

From the BINACLE assays for the activity determination of the toxin serotype BoNT/B, the following overall results were obtained in the collaborative study:

One remarkable result was that the signal intensities measured for the BoNT/B-containing sample solutions differed markedly between the participants. Only two out of ten laboratories obtained comparatively high toxin signals, which enabled them to sensitively detect even low toxin concentrations of less than 0.01 U/mL BoNT/B. The remaining eight participants, in contrast, obtained considerably lower toxin-induced signals, which made sensitive measurements difficult and, in several cases, also prevented valid RP calculations. But nevertheless, when focusing on the RP estimates resulting from all valid BoNT/B data sets obtained in the study, we found that these values showed only moderate variability between the laboratories. In fact, the degree of variability observed in the BoNT/B potency results was very similar to that seen in the BoNT/A BINACLE tests. This finding demonstrates that the BINACLE assay for BoNT/B is basically suitable for achieving reliable and reproducible potency measurements.

The RP estimates that we calculated based on the BINACLE assay results do not exactly reflect the unit specifications provided by the manufacturers of the respective BoNT products: According to the manufacturers’ specifications, the stock solutions of OnabotulinumtoxinA and IncobotulinumtoxinA used in the study each had an activity of 200 U/mL, and the stock solution of RimabotulinumtoxinB had an activity of 5,000 U/mL. Compared to the manufacturer’s specifications, the RP values that were calculated in our study for the two BoNT/A products were slightly higher (i.e. in the range of 230-330 U/mL). The RP values that we determined for the BoNT/B product, by contrast, were approximately 2- to 3-fold lower than the activity units stated by the manufacturer.

However, it is known that due to the lack of standardization of the *in vivo* potency tests for BoNT and also due to the fact that no internationally recognized standard toxins were available until recently, the activity units used by different manufacturers for their BoNT products are product-specific and non-equivalent [20;21]. Therefore, an absolute agreement between the activity units indicated by the toxin manufacturers for their respective pharmaceutical products and the potency estimates that were calculated relative to the research-grade toxin preparations which served as reference material in our study is not to be expected.

A root-cause analysis was performed to investigate the reasons for the remarkably high differences between the signal levels measured by the individual participants in the BoNT/B BINACLE assays. It revealed that the microplate washing conditions represent a relevant influencing factor: We observed that manual washing procedures tend to result in lower toxin signals in the BoNT/B BINACLE assay than procedures that use an automated washer. A key difference between manual and automated procedures may be the amount of washing buffer (PBS/0.05% Tween^®^ 20) remaining in the plate. It was previously reported that the proteolytic activity of tetanus neurotoxin, a toxin showing high structural and mechanistic similarities to BoNT/B, is negatively affected by traces of Tween^®^ 20 [19]. If BoNT/B exhibits a similar sensitivity to detergents, elevated amounts of washing buffer remaining in the plate after manual washing could potentially affect the synaptobrevin-cleaving activity of BoNT/B. In line with this assumption, follow-up experiments confirmed that the toxin signals obtained with manual washing can be markedly improved by using a detergent-free buffer in the final washing cycle before the synaptobrevin cleavage incubation.

However, even when applying such detergent-free washing steps, the signals obtained for BoNT/B using manual washing were still somewhat lower than those obtained using a washer. This suggests that other differences between the washing procedures, such as the contact time with the washing buffer, may also impact the BINACLE results.

Our findings illustrate that when implementing the BoNT/B BINACLE assay, the washing conditions require special attention. Using a microplate washer is generally recommended because automated procedures are more standardized and less susceptible to user-related variability than manual procedures. In cases where washing needs to be performed by hand, it may be beneficial to add cycles using Tween^®^ 20-free buffer prior to detergent-sensitive incubation steps.

For the BoNT/A BINACLE assay, in contrast, we did not find any evidence of increased sensitivity to the washing conditions.

The fundamental suitability of BINACLE assays for use in the batch testing of pharmaceutical products is substantiated by the results of a former collaborative study organized by the European Directorate for the Quality of Medicines & HealthCare [14]. The data obtained in this study, which aimed to characterize the applicability of BINACLE assays for detecting residual tetanus neurotoxin during vaccine production, confirmed that BINACLE assays allow reproducible and sensitive measurements of toxin activity. Based on these results, the BINACLE assay for tetanus neurotoxin has recently been included into the European Pharmacopoeia as an alternative method for testing the safety of tetanus toxoids [22;23].

Before BINACLE assays can be used as alternative method for the activity or potency determination of pharmaceutical BoNT preparations, a laboratory- and product-specific validation is strictly needed. A crucial aspect of such validation procedures is to carefully check whether the results of the assay closely reflect *in vivo* activity and whether they reliably quantify losses in activity caused by factors such as heat stress or prolonged storage.

Another crucial point for the routine applicability of the method is the long-term availability of suitable reagents. At present, not all specific reagents needed for the BINACLE assays are commercially available as regular catalogue products. For example, for detection of the cleaved substrate proteins SNAP-25 and synaptobrevin, polyclonal rabbit antibodies produced by contract manufacturing are currently used. For future use of the method, a switch to monoclonal or recombinant antibodies would be preferable for reasons of animal protection, as well as to reduce batch-to-batch reagent variability. Investigations into this matter are currently underway.

Reference toxins are another important factor: Well-characterized toxin preparations should be included in each BINACLE test as positive controls to verify that the test has been performed correctly and that the results are valid. Additionally, they serve as a reference point for calculating relative potency. Ideally, certified standard toxins should be used for this purpose. At the time when our collaborative study was conducted, no internationally recognized standard material for BoNT was available. We therefore used commercial, research-grade toxin preparations as an internal reference. Recently, certified BoNT standard preparations have become available [24]. Such standardized materials are expected to be valuable tools for future studies, as they facilitate the reliable assignment of relative potency values that can be reproduced in different laboratories.

Taken together, the collaborative study has demonstrated that for samples containing BoNT/B, an application of the BINACLE assay for activity measurements is possible, and reproducible RP calculations can be achieved. Furthermore, the study revealed useful information regarding the sensitivity of BoNT/B to detergents, which provided valuable insights into how the protocol can be modified to enhance the robustness of the BoNT/B BINACLE assay. However, it should be noted that, following the withdrawal of Neurobloc^®^’s marketing authorization in 2022, no BoNT/B-based products are currently available on the European market. Consequently, *in vitro* methods for BoNT/B activity measurement have a limited relevance for pharmaceutical applications. Nevertheless, the BoNT/B BINACLE assay represents a valuable tool for research applications.

BoNT/A, by contrast, is by far the most relevant toxin serotype for pharmaceutical purposes, and numerous products containing this toxin serotype are approved for various applications in clinical and aesthetic medicine. Methods that allow reliable measurements of BoNT/A activity are therefore of great importance. The results of our study indicate that the BINACLE assay for determining the activity of BoNT/A is robust and allows reproducible relative potency determinations of pharmaceutical BoNT/A products. Furthermore, the data demonstrate that the method can be straightforwardly transferred to new users with no prior BINACLE experience. Consequently, we conclude that the BoNT/A BINACLE assay represents a suitable method for *in vitro* activity measurements, fulfilling the basic requirements for use as an alternative to the mouse bioassay.

In summary, our collaborative study confirms the utility of the BINACLE assay for determining the potency of BoNT products, offering a promising basis for its future applications in this field.

## Acknowledgements

We thank all participants for their valuable contribution to the study. We also wish to express our sincere gratitude to Beate Krämer and Emina Wild for their substantial contributions to the development and optimization of the BINACLE test principle. Without their efforts, the collaborative study would not have been possible.

The project was funded by the German Federal Ministry of Education and Research (BMBF projects no. 031A210 and 161L0148) and the German Research Foundation (DFG, Ursula M. Händel Animal Welfare Prize 2016). The funding institutions did not influence the study design, the analysis and interpretation of the data, and the writing of the report.

## Author contributions

Study design: BK, HBN and KMH. Preparation of the experimental protocol for the study: UB, JK, BK and HBN. Pre-qualification and shipment of reagents: JK, UB and HBN. Collection and review of data: BK and HBN. Statistical data analysis: KMH. Evaluation of the results and drafting of the article: HBN and BK. Critical revision and editing of the manuscript: all authors.

## Participants of the collaborative study (listed in alphabetical order by the name of the respective company or institute)

L. Töllner, CROMA-PHARMA GmbH, Austria

S. Jorajuria, G. Cozic, European Directorate for the Quality of Medicines & HealthCare (EDQM), Council of Europe, France

M. Wall, D. Lam, Health Canada, Canada

M. Kochte, M. Malkowsky, IDT Biologika GmbH, Germany

R. Zichel, E. Dor, O. Rosen, Israel Institute for Biological Research (IIBR), Israel

R. Tierney, P. Stickings, Y. Liu, National Institute for Biological Standards and Control (NIBSC), United Kingdom

H. Behrensdorf-Nicol, U. Bonifas, J. Klimek, K.-M. Hanschmann, B. Kegel, Paul-Ehrlich-Institut, Germany

D. Stern, B. Dorner, O. Shatohina, Robert Koch Institut, Germany

A. Rummel, N. Krez, toxologics GmbH, Germany

F. Neuschäfer-Rube, Universität Potsdam, Institut für Ernährungswissenschaft, Germany

## Abbreviations

ANOVA: analysis of variance
AU: absorbance units
BINACLE: binding and cleavage
BoNT: botulinum neurotoxin
BSA: bovine serum albumin
PEI: Paul-Ehrlich-Institut
PBS: phosphate-buffered saline
PIPES: 1,4-piperazinediethanesulfonic acid
RP: relative potency
SD: standard deviation
SNAP-25: synaptosomal-associated protein-25
SV2C: synaptic vesicle glycoprotein 2C
TCEP: tris(2-carboxyethyl)phosphine hydrochloride
TMAO: trimethylamine N-oxide dihydrate
TMB: 3,3’,5,5’-tetramethyl benzidine

